# Increasing the predictive accuracy of the Resistance Gene Identifier by evaluating antimicrobial resistance gene over- and underprediction

**DOI:** 10.64898/2025.12.11.693720

**Authors:** Karyn M. Mukiri, Brian P. Alcock, Amogelang R. Raphenya, Andrew G. McArthur

## Abstract

Computational biology is paving the way towards accessible antimicrobial resistance (AMR) gene (ARG) detection methods to complement canonical gold-standard phenotypic diagnostics for the purposes of antimicrobial surveillance. However, obtaining an accurate depiction of phenotypic resistance through genotypic methods requires addressing potential over- and underprediction of ARGs. This study assessed the manual curation accuracy of bit-score cutoff values associated with bioinformatic models within the Comprehensive Antibiotic Resistance Database (CARD) and the subsequent effects of erroneous cutoff curation, leading to potential Type I and Type II error, on its *in silico* resistome prediction tool, the Resistance Gene Identifier (RGI). CARD models rarely overpredicted (5 of 3,900 models with >5% false positive rates) but somewhat underpredicted (739 of 3,900 models with >5% false negative rates) resistance-associated sequences and mutations, emphasizing RGI’s conservative prediction algorithms. Isolating curation inaccuracies by AMR gene family, efflux-related families were the main contributors to overprediction (likely due to human curation error), while underprediction was primarily due to beta-lactamase families, the latter finding highlighting systemic curation deficiencies.

## INTRODUCTION

Antimicrobial resistance (AMR), one of the top ten threats to global health (1), places great strain on healthcare systems worldwide. In 2019 alone, it was estimated that of the 4.95 million deaths associated with AMR, ∼25.66% (1.27 million) deaths were directly attributable to resistance (2), and, without further intervention, global deaths directly attributable to AMR could reach 39 million between 2025 and 2050 (3). Due to factors such as clinical overuse of antimicrobial agents, use of substandard antimicrobials, as well as their overuse in both animal husbandry and agriculture (4–6), the worldwide burden of resistance has seen a steady increase since the first reported instance of bacterial resistance in 1942 (7). This widespread misuse has led to the decreased efficacy of key clinical antibiotics, including our antibiotics of last resort, as bacteria evolve to combat these mechanisms (8, 9). Moreover, the evolution of novel bacterial resistance mechanisms makes the task of engineering new and novel therapeutics extremely difficult; this has left our global clinical pipeline in a profoundly compromised state (10).

With DNA sequencing technology making monumental strides in its development, outpacing Moore’s Law in 2008 (11), computational biology is paving the way towards accessible resistance gene detection methods to complement canonical gold-standard phenotypic diagnostics. Proceeding this was a wave of big data, which called for a proactive approach towards standardizing and organizing nomenclature and knowledge of AMR mechanisms through the use of curated databases (12). A handful of core databases that work to direct the community via their meticulous curation methods are the National Center for Biotechnology Information (NCBI) Pathogen Detection Reference Gene Catalog (13), ResFinder (14), and the Comprehensive Antibiotic Resistance Database (CARD) (15).

CARD sits amongst the aforementioned resources as a manually curated, ontology-centric database containing 8,582 ontology terms and 6,442 reference sequences (version 4.0.1) through which it performs *in silico* resistome annotation using its in-house Resistance Gene Identifier annotation software (RGI; https://github.com/arpcard/rgi). Essential for predicting AMR genes (ARGs) and mutations from newly acquired genomic sequences is CARD’s Model Ontology (MO), a controlled vocabulary (16) that describes the parameters used in the detection of AMR determinants across a widely diverse range of bacterial genomes (17). Overall, CARD’s ontologies give its annotation software the context required to better describe and annotate bacterial resistomes, i.e., collections of all possible antibiotic resistance genes within bacterial chromosomes and plasmids (18). Additionally, CARD includes the Resistomes and Variants (CARD-R) dataset, a collection of isolate-based resistome predictions and associated ARG prevalence data.

CARD includes eight bioinformatic detection model types for ARG prediction by RGI (e.g., the protein homolog model, a presence/absence search) which are wholly reliant on the MO, in addition to the Antibiotic Resistance Ontology (ARO), for informative genome annotation. For example, consider CTX-M-15 (19), a beta-lactamase found in the *Enterobacteriaceae* family, encoded by a protein homolog model using a reference protein sequence (sourced from NCBI) and a curated BLASTP bit-score cutoff (a sequence alignment based similarity detection parameter). If detected by RGI, the ARO would then define the AMR gene family (CTX-M beta-lactamase), drug class (penam, cephalosporin), and related AMR information. However, to date, only four of eight total AMR detection model types curated in CARD are utilized by CARD’s RGI software, three of which do not account for all possible detection parameters included in CARD (e.g., specific kinds of mutations such as indels) and are, thus, underpredicting AMR determinants when annotating genomes or metagenomes. Furthermore, the most frequently used model type (protein homolog) is highly dependent on the accurate hand curation of the bit-score cutoff sequence homology detection parameter, which may require further fine-tuning to increase confidence in predictive gene predictions across several models.

RGI has three analytical branches: *main* (for annotation of genome or plasmid sequences, genome assemblies, metagenomic assemblies, or proteomes), *bwt* (for annotation of metagenomic sequencing reads), and *k-mers* (for prediction of pathogen-of-origin for ARGs). RGI *main* uses two pairwise local sequence alignment tools, BLAST (20) and DIAMOND (21), to predict AMR determinants homologous to reference sequences in CARD when analyzing genomic data and is the focus of this study. Reliability of metagenomic sequencing reads by RGI *bwt* will be the focus of a separate study. Overall, RGI *main* compares user-provided sequences to the reference sequences within CARD, such that predicted annotations are likely to be functional ARGs. To achieve this, BLAST or DIAMOND calculate a statistical value, bit-score, for each annotation, which approximates our confidence in the query-to-reference pairwise alignment accuracy. The bit-scores of individual annotations are then compared to the CARD reference bit-score cutoffs set by CARD biocurators; pairwise BLAST or DIAMOND alignments with equal or greater bit-scores than the curated cutoffs are assumed to be a functional variant of the reference ARG curated in CARD. CARD biocurators determine bit-score cutoffs by performing all-against-all BLASTP (protein versus protein) of the reference sequences in CARD to determine bit-score values that differentiate pairwise alignments within AMR gene families from pairwise alignments among families, e.g., all CTX-M beta-lactamase sequences would have bit-score values above the cutoff with alignment to other CTX-M beta-lactamases but below the cutoff for alignment to FONA, SFO, OXY, and RAHN beta-lactamases. More details on the hand curation of bit-score cutoffs can be found in the RGI documentation (https://github.com/arpcard/rgi).

Using these curated bit-score cutoffs, RGI *main* annotations are separated into three categories based on their level of similarity to a CARD reference sequence: RGI Perfect, RGI Strict, and RGI Loose. RGI *main* defines Perfect annotations as gene calls with 100% amino acid sequence similarity to a CARD reference sequence, while Strict annotations are not exact matches but are highly similar and likely functional based on CARD bit-score cutoffs (i.e., BLAST or DIAMOND alignment bit-score equals or surpasses CARD bit-score cutoff). Loose annotations occur when the BLAST or DIAMOND alignment bit-score is less than CARD bit-score cutoff and are low similarity predictions that may or not reflect ARGs. Combined with phenotypic screening, the Loose algorithm allows researchers to home in on new AMR genes (15).

To ensure confidence in RGI *main* ARG annotations, manual biocuration and validation of ARO terminologies and associated data, including the parameter values for each AMR detection model, is imperative. Inaccurate curation may leave RGI *main* gene predictions and annotations liable to both Type I and Type II error. RGI Perfect annotations are not affected by this as they are defined by an exact amino acid match to a CARD reference sequence; however, the differentiation between RGI Strict and RGI Loose annotations is wholly dependent upon manually curated CARD bit-score cutoff values. Cutoff values set too low will lead to an increased number of spurious, false positive Strict annotations, while cutoffs set too high will result in false negative Loose annotations. Thus, in this study, we detail an exhaustive investigation into CARD bit-score cutoff curation and its subsequent effects on RGI *main* ARG annotation.

## MATERIALS AND METHODS

### Accounting for conservative amino acid substitutions

To find specific instances of erroneous bit-score cutoff curation in CARD, it was necessary to look beyond RGI’s sole reliance on bit-score and percent identity (the number of exact matches within a pairwise alignment). The RGI v5.2.1 base code was modified to obtain an additional alignment statistic not included in standard RGI output: percent positive. Percent positive is a similarity statistic that considers the number of exact matches within an alignment, whilst additionally factoring in conservative substitutions (e.g., leucine to isoleucine) (22, 23); thus accounting for the physico-chemical properties of amino acids lends a more all-encompassing similarity metric.

### Assessment data

CARD’s Resistomes & Variants sequence data (CARD-R) reflects computer-generated resistome predictions for important pathogens and includes ARG sequence variants beyond those reported in the scientific literature. CARD-R v3.1.0 contained data from 263 pathogens, 16,719 chromosomes, 2,675 genomic islands, 33,860 plasmids, 136,704 WGS assemblies, and 285,146 ARG alleles, based on both Perfect and Strict RGI *main* annotations, with the latter scoring above the curated bit-score cutoff and thus possibly containing false positive annotations. Like all CARD-R releases, RGI Loose annotations were excluded, given their need for further experimental validation, yet these may include false negative annotations. To accurately evaluate curated bit-score cutoff accuracy across the entirety of CARD, we re-analyzed CARD-R v3.1.0 using our modified version of RGI and CARD v3.2.1. Our assessment dataset (mCARD-R) included 150,177 sequences of RGI Perfect, Strict, and Loose annotations along with their bit-score, percent identity, and percent positive values. The RGI decision criteria were not changed (i.e., bit-score cutoffs to differentiate RGI Strict and Loose) as percent identity and percent positive were reserved for secondary interpretation (outlined below). DIAMOND v0.8.36 alignment was used as opposed to the RGI default (BLAST) as the DIAMOND algorithm had a speed advantage due to its use of spaced seeds (segments of longer seeds) as opposed to single, consecutive seeds (21), as well as to preserve comparability with the original CARD-R annotations, which were also generated using DIAMOND. RGI’s nudged annotations option (≥95% identity Loose annotations automatically annotated as Strict) was also excluded as this would introduce an unnecessary level of ambiguity between models. Finally, the percent identity and percent positive statistics used in our analyses were generated by DIAMOND using the BLOSUM62 substitution matrix to represent conservative amino acid substitutions.

### Assessment criteria

Taking advantage of both percent positive and the percent length of the CARD reference sequence that the alignment of the query protein encompasses, RGI annotations were secondarily interpreted. This allowed for the creation of two new annotation subcategories – False Strict (FS) and False Loose (FL) – to isolate distinct instances of RGI over- and underprediction, respectively. Our assumption is that FS (<70% percent positive alignments) are proteins unrelated to AMR yet annotated as RGI Strict annotations based on RGI’s default use of bit-score. Conversely, FL (≥70% positive alignments; ≥70% of the reference sequence length) are assumed to be annotations that are likely functional ARGs yet annotated as RGI Loose annotations based on RGI’s default use of bit-score. The selection of 70% percent positive to differentiate accepted from False annotations is discussed below.

### Phylogenetic trees

RGI predicted protein sequences of target AMR gene families were selected from the mCARD-R dataset and consolidated into a merged FASTA file for multiple sequence alignment preparation, along with CARD reference protein sequences for individual models. A secondary file with selected metadata (model/ARO name and bit-score cutoff) for each RGI annotation was also prepared. Multiple sequence alignments were created with Augur v24.3.0 and trimmed using trimAl’s automated1 method v1.4.rev15 (build[2013-12-17]). Phylogenetic trees were constructed using IQ-TREE’s v2.3.0 default ModelFinder Plus (MFP) with 1000 bootstrap replicates. The final trees were midpoint rooted and visualized using the R package ggtree v3.10.1.

## RESULTS

### Assessment data

The generated assessment data (mCARD-R) encompassed 255 pathogens and generated 56,708,241 Perfect (1,073,204), Strict (2,584,487), and Loose (53,050,550) annotations based on the unmodified RGI decision criteria (i.e., bit-score). In total, mCARD-R included annotations for 3,900 of CARD v3.2.1’s 4,788 detection models. The remaining 938 detection models reflect ARGs not present in the original CARD-R data, e.g., very rare beta-lactamases.

### RGI over- and underprediction

Based on our assessment criteria, the range of Perfect, Strict, and Loose annotations generated is readily distinguishable (Figure 1). Some models, such as TEM-1, generated very few, if any, False annotations. Compare this to efflux protein models such as adeF and YojI, which frequently generate False Strict and False Loose annotations, respectively (Figure 1). We also examined this across all of CARD by viewing the rate at which either false positives (Figure 2A) or false negatives (Figure 2B) are generated by CARD’s bioinformatic models. Overall, CARD models underpredicted some resistance-associated sequences and mutations (i.e., 739 of 3,900 models with >5% False Loose rates), yet there was very little overprediction of annotations (i.e., only 5 of 3,900 models with >5% False Strict rates). Since it is not feasible to address all instances of underprediction due to the required biocuration burden, we here primarily focus on annotation models that generated these annotations at frequencies of 20% or higher. This subset further emphasized RGI’s conservative prediction algorithms, as only 0.13% of models produce FS annotations, while 11.23% produce FL annotations at these rates.

**Figure 1.**
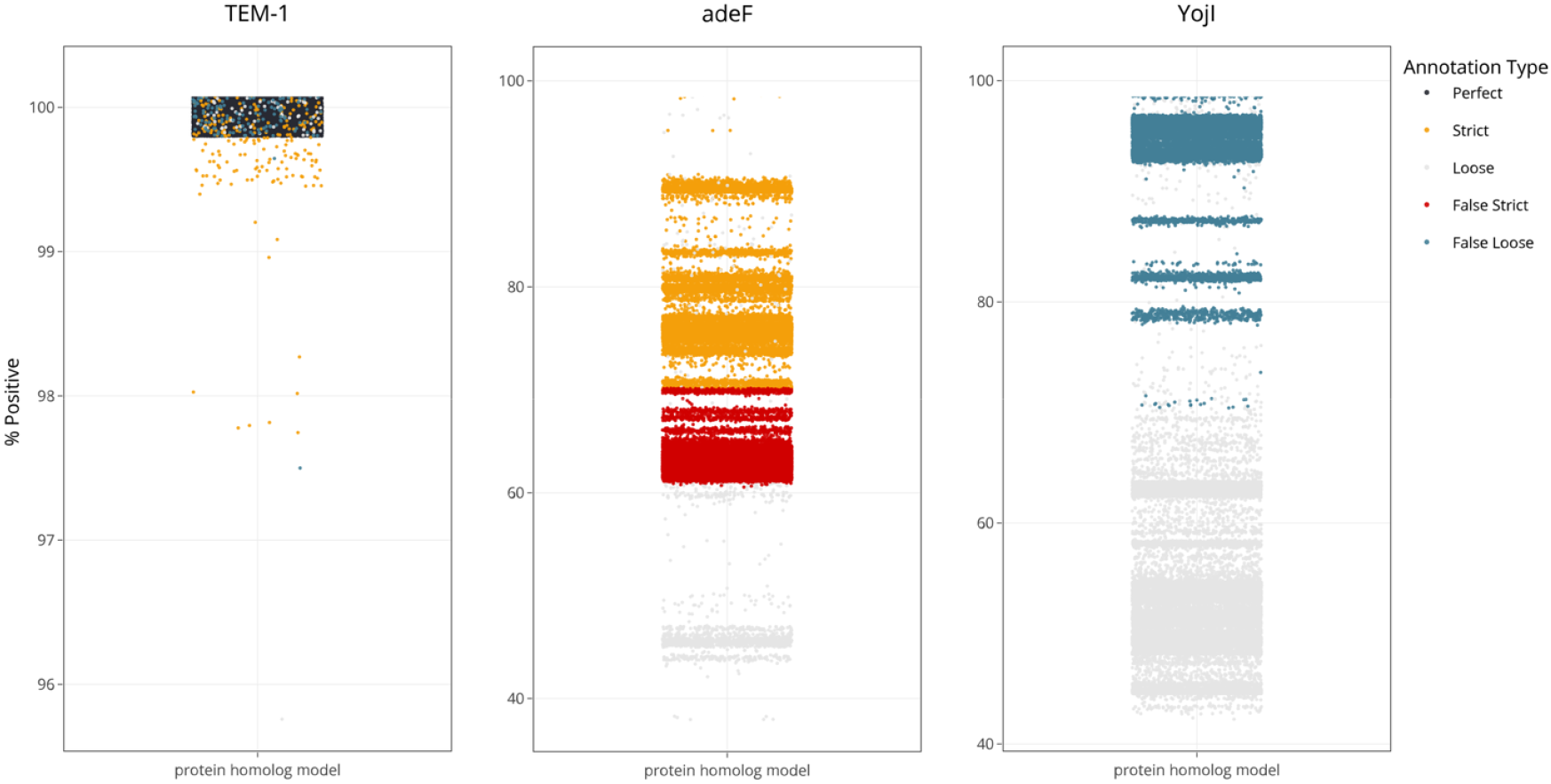
False Strict and False Loose RGI annotations for three example resistance genes. The percent positive statistic of each annotation is plotted along the y-axis. Annotations are categorized as Perfect, Strict, False Strict, Loose, and False Loose based on our assessment criteria. TEM-1 is an example of a well-performing annotation model (0% False Strict, 0.75% False Loose), while adeF (48.58% False Strict, 0% False Loose) and YojI (0% False Strict, 31.92% False Loose) showcase models with annotation challenges that may be powered by incorrectly curated bit-score cutoffs.

**Figure 2.**
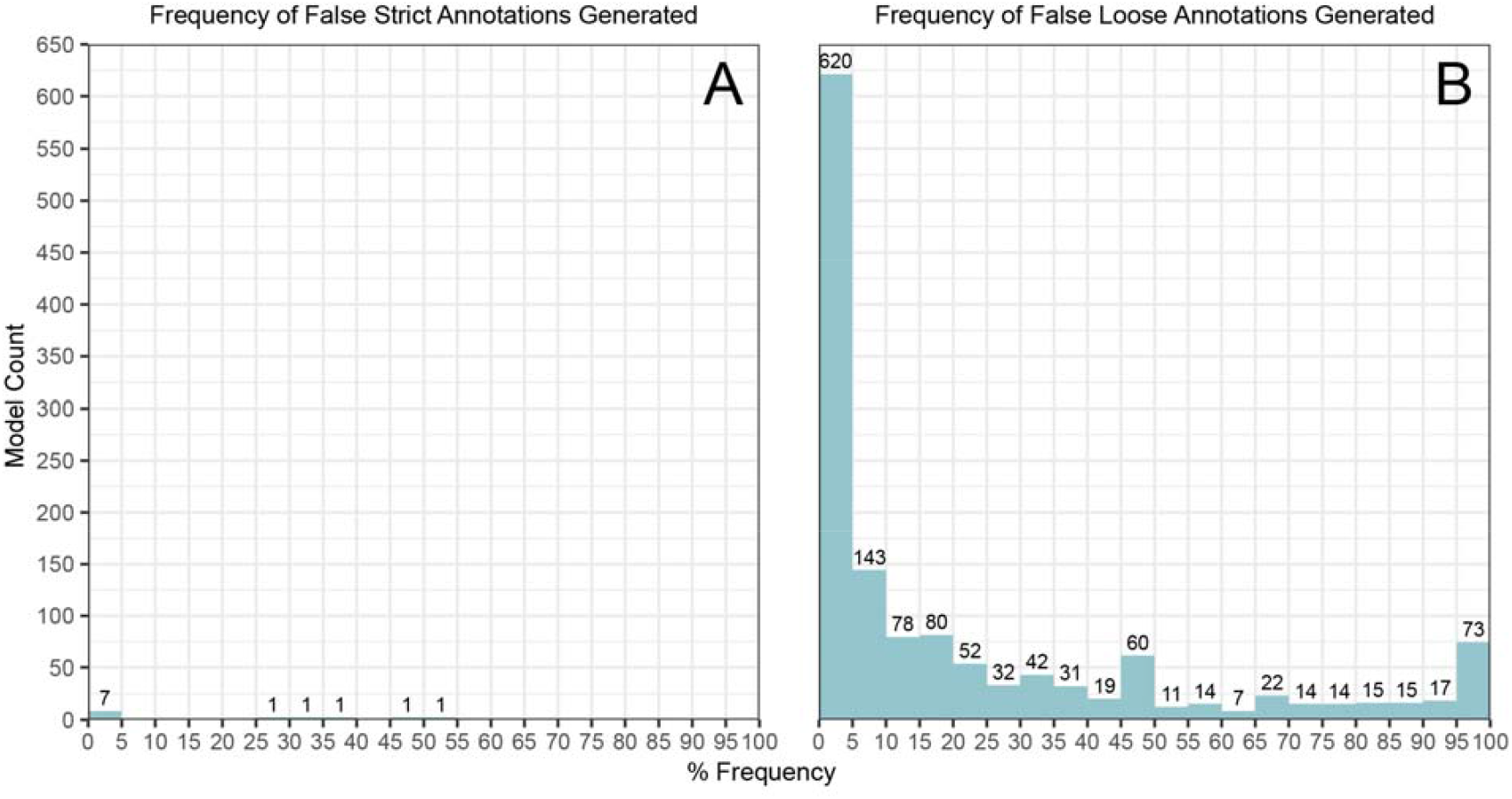
Frequency of False Strict and False Loose annotations in the entire mCARD-R assessment data. Each frequency bin contains a distinct number of ARG annotation models and the bins are categorized by the frequency at which these models generate either False Strict (A) or False Loose (B) annotations. The total number of models is presented at the top of each bin. For example, the adeF model outlined in Figure 1 is in the 45-45% False Strict bin and 0-5% False Loose bin.

### AMR gene families

Genes and mutations within CARD’s Antibiotic Resistance Ontology (ARO) are associated with three major ontological categories: AMR gene family, drug class, and resistance mechanism. To tease out potential patterns of erroneous bit-score cutoff curation, we shifted our focus towards AMR gene families instead of individual genes. Setting aside gene families with 10 or less genes, nine families included generation of False Strict annotations, while 29 families included generation of False Loose annotations. For context, CARD v3.2.1 included 444 AMR Gene Families total. Figure 3A outlines models largely responsible for the rare cases of possible ARG overprediction, of which four of five annotation models were for antibiotic efflux proteins. Due to the low number of annotation models involved, we assume that the overprediction of ARGs by RGI is the result of human error (i.e., inexperienced curators, bit-score cutoff curation rules improperly followed). Figure 3B illustrates the overwhelming role beta-lactamases play in ARG annotation underprediction, as they constitute eight of the top 10 underperforming ARG families. Given the severity of these numbers, we hypothesized that this level of underprediction was the result of a deeper issue.

**Figure 3.**
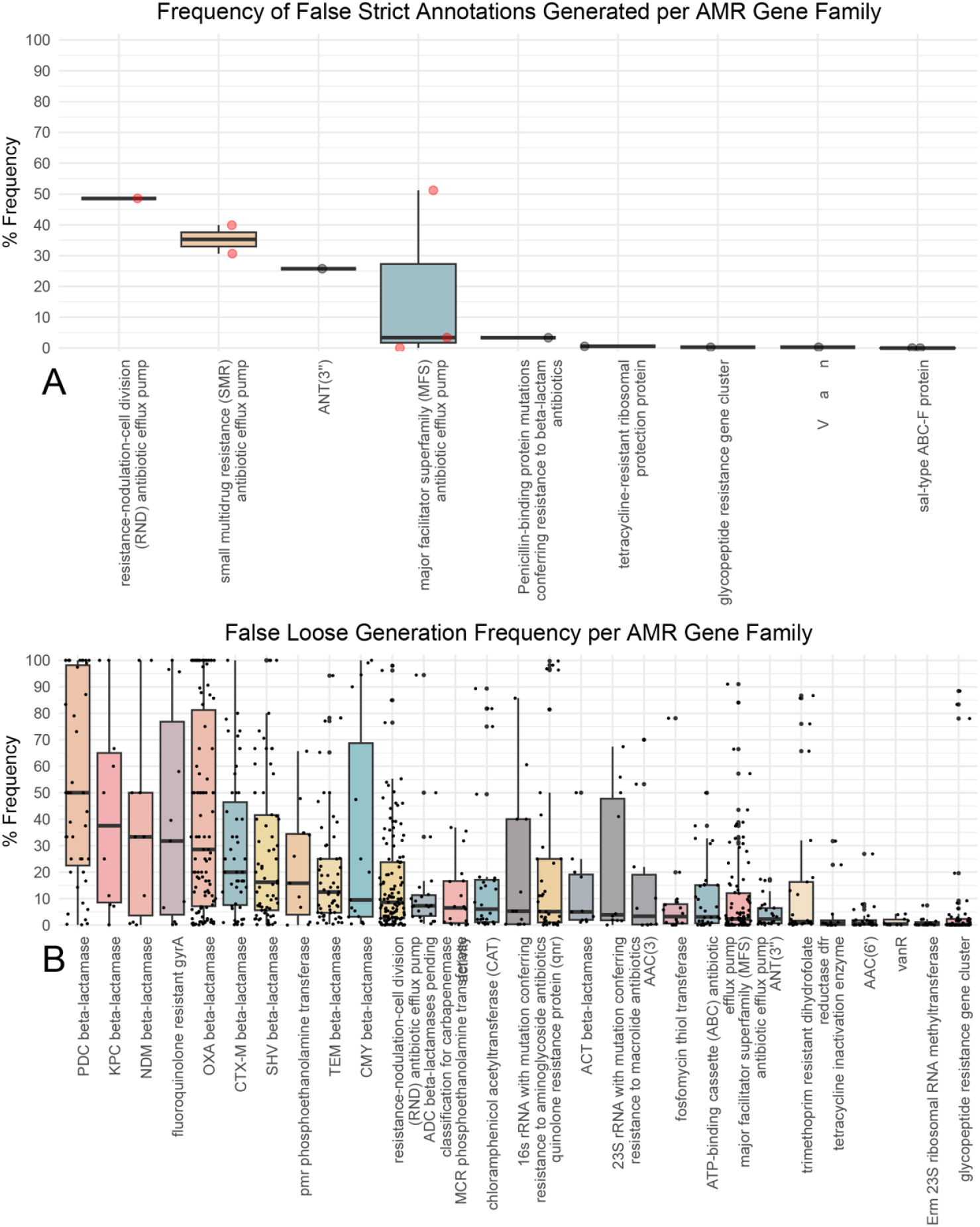
Isolating patterns of bit-score cutoff erroneous curation by investigating the frequency of False Strict (A) and False Loose (B) annotations for AMR gene families. Families consisting of 10 or fewer genes have been excluded. Each dot represents an individual ARG annotation model, with efflux proteins highlighted in red. AMR gene families (x-axis) are plotted in descending order of their median false annotation rate.

### Phylogenetic context

We examined curation practices around beta-lactamases as a whole, comparing CARD to NCBI’s Pathogen Detection Reference Gene Catalog, the latter being the main curation body responsible for cataloguing newly discovered beta-lactamases (24). New alleles are accepted and named solely based on their amino acid sequence, and every new variant is archived to support AMR surveillance, regardless of the presence of an experimentally validated resistance phenotype. This has led to an influx of new variants, which CARD, at times, bulk imports from NCBI into its own database. As variants sometimes only differ by 1-2 amino acids, it is often not feasible for CARD curators to assign each individual beta-lactamase protein a unique bit-score cutoff, considering the sheer number of new incoming variants. To combat this, curators sometimes assign a blanket bit-score cutoff to incoming batches of beta-lactamases based on past curation. Yet, gene subfamilies have also been identified for some beta-lactamase families, such as those within the OXA family (25), and these could guide more accurate bit-score curation efforts. To explore potential sub-familial relationships within other beta-lactamase families, phylogenetic trees were constructed for the eight beta-lactamase families generating high rates of FL (sequences extracted from CARD version 3.2.6). For the OXA beta-lactamases, we observed the same OXA subfamily structure as found in the literature (Fig. 4A), with potentially new subfamily clades identified. Yet, Figure 4 also highlights how bit-score curation is often incongruent with phylogenetic structure (OXA, KPC, NDM, SHV, PDC, and CMY beta-lactamases) or ignores phylogenetic structure entirely (TEM & CTX-M beta-lactamases). We hypothesize that curation of bit-scores for beta-lactamases without considering phylogenetic relationships among variants has led to high rates of FL for the large beta-lactamase families in Figure 4.

**Figure 4.**
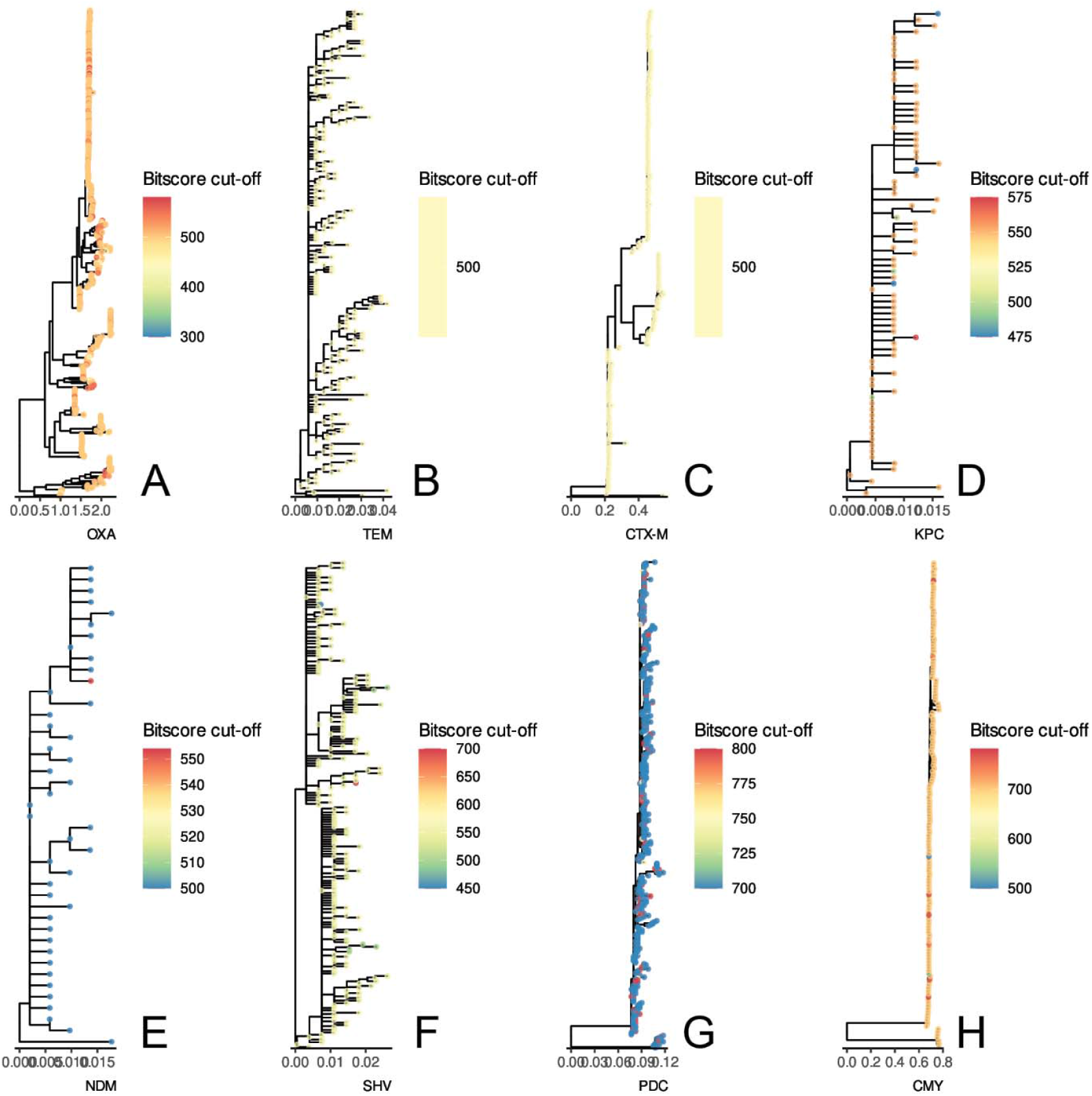
Highlighting subfamily organization and bit-score curation discrepancies for eight beta-lactamase gene families with high rates of False Loose annotations through phylogenetic analysis: OXA (A), TEM (B), CTX-M (C), KPC (D), NDM (E), SHV (F), PDC (G), CMY (H). Trees are plotted with the number of protein substitutions per site on the x-axis and nodes are coloured by their respective, curated CARD bit-score cutoffs.

In contrast, the MCRs (a family of phosphoethanolamine transferases conferring resistance to colistin), whose nomenclatural system was established following the schema set in place for beta-lactamases (e.g., MCR-1.1, MCR-2.1, etc.), did not produce FL annotations at high rates. The MCR naming schema runs in a similar fashion, however, subfamilies are created and populated based on amino acid sequence variance and common ancestry (26). Members within a subfamily are usually 1-2 amino acids different from the wild type/major family member, while entirely new families may be lower than ∼88% identical to one another. CARD’s curation of bit-scores follows these MCR subfamilies.

## DISCUSSION

The reliability of CARD’s RGI software is dependent on minimizing both the under- and overprediction of ARGs by ensuring the maximal curation accuracy (i.e., overall correctness) of CARD’s bioinformatic models. For many, overprediction is the main concern, as false positive results can affect downstream analyses; even though RGI only predicts the *in silico* resistome (and not phenotype), users do not want to be misled on potential phenotypic susceptibility. While RGI Perfect annotations are self-evident (i.e., identifying proteins 100% identical to curated references), our systematic evaluation has shown that only five of 3,900 of CARD’s annotation models are frequently overpredicting RGI Strict annotations (>5% error rate). We thus conclude that CARD’s conservative approach to curating bit-score cutoffs results in high RGI Strict precision, i.e., annotations generated overwhelmingly identify resistance genes. Notably, CARD’s efflux models are notoriously difficult to curate due to a general inability to distinguish between AMR and non-AMR homologs and poor surrounding experimental evidence, as they are difficult to manipulate. Conversely, CARD annotation models which underpredict RGI Strict annotations pose a systemic curation challenge; the prediction of potential antibiotic resistance is being undermined by poor recall. Yet, only 739 of 3,900 (18.9%) of CARD’s annotation models have False Loose rates of greater than 20%. At a practical level, distant variants annotated as RGI Strict (or False Loose) should always be interpreted with some caution, given their divergence and lack of functional data. We note that none of the above obviates the basic assumptions of CARD and RGI: (i) prediction of resistance genotype does not guarantee resistance phenotype, (ii) based on bit-score cutoffs, RGI Strict annotations are likely functional AMR genes, but expression and phenotypic screening is required for certainty, and (iii) RGI Loose annotations can include new, emergent threats and more distant homologs of AMR genes, but will also catalog homologous sequences and spurious partial matches that may not have a role in AMR.

The 70% positive alignment score we selected for our assessments of RGI Strict and Loose performance was an exploratory starting point to tackle egregious errors in bit-score cutoff curation. We cannot point to theoretical or experimental justifications, as there are different variables that could factor into how cutoffs are set. However, in a study by Berglund *et al*., 76 putative beta-lactamases were identified using Hidden Markov Models (27). Within these data, five genes (G33, G37, G58, G70, and G71) conferred resistance to imipenem, yet were classified as RGI Loose; our analysis reclassified them as False Loose. Increasing the percent positive cutoff alters our results drastically (Supplementary Figure 1), as changing it in either direction determines stringency and either increases or decreases the flagging of False annotations. Additionally, similar to CARD’s bit-score cutoff curation, individual AMR gene families may benefit from specifically assigned percent positive cutoff values (Supplementary Figure 2), depending on how many models populate the family and how well annotated the family is as a whole. Certainly, a single percent positive cutoff across the diversity of proteins involved in AMR is hard to justify. This reinforces CARD’s core principle: dedication to human examination of data and hand biocuration of sequences and models. In this, our 70% positive cutoff is leading CARD’s biocurators to genes and gene families requiring closer inspection as well as spurring experimental validation by the AMR research community.

The large number of possibly underpredicting beta-lactamase models calls for a shift in CARD’s curation practices. While blanket bit-score cutoffs are convenient due to the sheer number of beta-lactamases that are regularly imported into CARD, it may be beneficial to instead assign blanket cutoffs for individual subfamilies. Our phylogenetic investigations have revealed needed improvements in CARD’s curation practices. As of CARD version 3.2.9, CARD has incorporated beta-lactamase subfamily curation to bolster bit-score curation practices and introduce more order within each individual gene family. Figure 3 indicates this should expand to other resistance mechanisms and that phylogenetic assessments should become an essential part of ARG nomenclature practices.

## CONCLUSIONS

Addressing both the over- and underprediction of ARGs by RGI brings us closer to a more complete picture of phenotypic resistance, and if the objective of ongoing curation for surveillance purposes is to be realized, our software needs to be able to rely on accurately curated data. Yet, if we reorient the 56,708,241 annotations provided by our mCARD-R assessment in terms of a confusion matrix with RGI Perfect, Strict, and Loose annotations as True Positives, False Strict as False Positives, and False Loose as False Negatives, we find RGI has 0% error rate, 100% accuracy, 100% precision, 99.96% recall, 100% specificity, and overall F1 score of 99.98% across 3,900 of CARD v3.2.1’s 4,788 detection models (values rounded to second decimal place, Supplementary Figure 3). This illustrates that while a small number of CARD’s detection models struggle with false predictions, based on our 70% percent positive cutoff, CARD’s overall curation strategy and RGI’s algorithms are highly reliable.

## SOFTWARE & DATA AVAILABILITY

Version-controlled copies of the Resistance Gene Identifier (RGI) are available at the CARD GitHub repository: https://github.com/arpcard/rgi.

## SUPPLEMENTARY DATA

Supplementary Data are available in association with this manuscript at *bioRxiv*.

## ACKNOWLEDGMENTS

Computational support was provided by the McMaster Faculty of Health Sciences Advanced Computing Facility, supplemented by hardware donations and loans from Cisco Systems Canada, Hewlett Packard Enterprise, and Pure Storage. AGM holds McMaster’s David Braley Chair in Computational Biology, generously supported by the family of the late Mr. David Braley. Feedback on this research was provided by Drs. Lori Burrows and Brian Coombes.

## AUTHOR CONTRIBUTORS

KMM: conceptualization, methodology, investigation, software, data curation, formal analysis, writing – original draft, writing – review and editing. BPA: data curation, supervision, writing – review and editing. ARR: software, writing – review and editing. AGM: conceptualization, funding acquisition, project administration, supervision, writing – review and editing.

