## Supplementary Figures 1-3 for "Increasing the predictive accuracy of the Resistance Gene Identifier by evaluating antimicrobial resistance gene over- and underprediction"

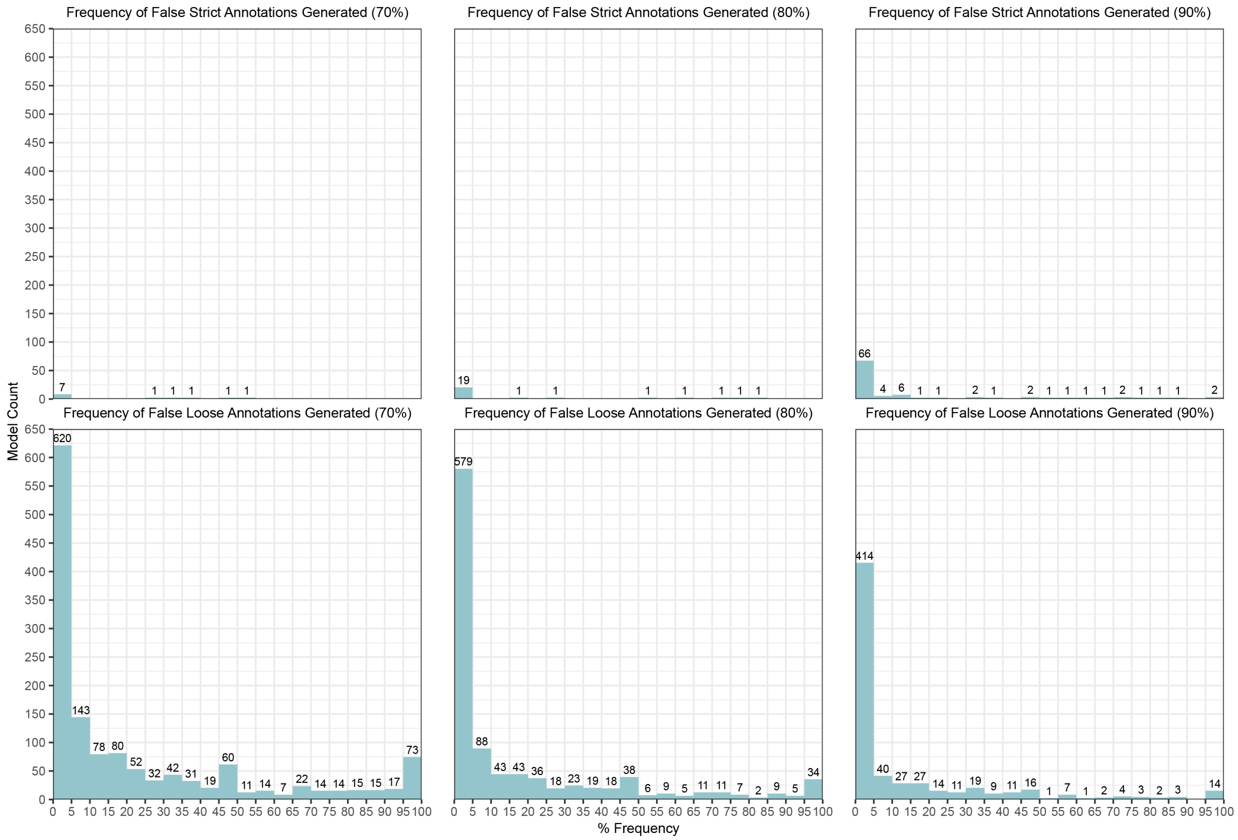


**Supplementary Figure 1**. The effects of increasing False annotation rate cutoffs on both False annotation types. Increasing False annotation filtering cutoffs are shown in increasing order from left to right (top: False Strict; bottom: False Loose). Each frequency bin contains a distinct number of models, e.g. ARGs. The bins are categorized by the percent frequency at which either False Strict (top) or False Loose (bottom) annotations are generated by RGI *main*.

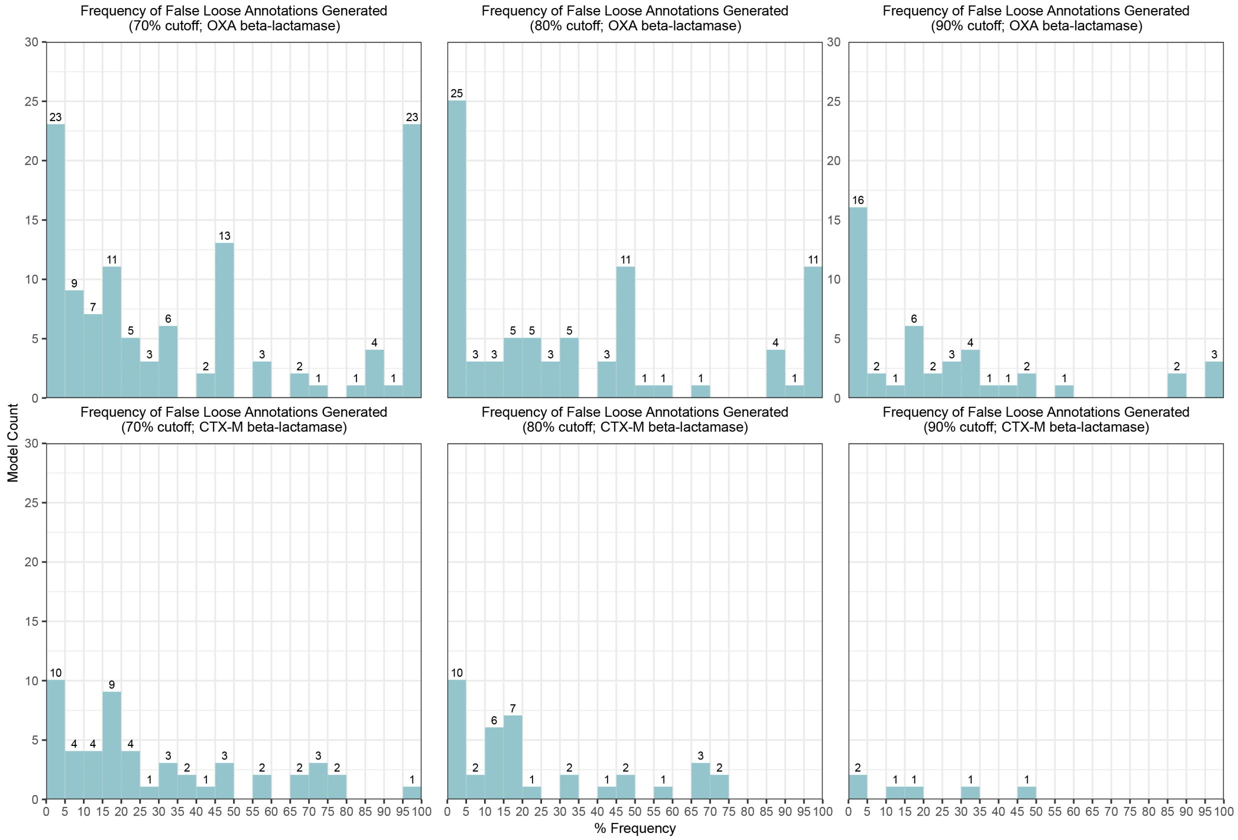


**Supplementary Figure 2**. The effects of different False annotation rate cutoffs on a model-by-model basis for OXA and CTX-M beta-lactamases. Increasing False Loose annotation flagging cutoffs are shown in increasing order from left to right (top: OXA beta-lactamases; bottom: CTX-M beta-lactamases). Each frequency bin contains a distinct number of models, e.g. ARGs. The bins are categorized by the percent frequency at which False Loose annotations are generated by RGI *main*.

**56,708,241 Perfect, Strict, or Loose annotations from 3,900 CARD models**

| Result | Category | Annotations |
|---|---|---|
| **Perfect** | TP | 1073204 |
| **Strict** | TP | 2584475 |
| **Loose** | TN | 53049191 |
| **False Strict** | FP | 12 |
| **False Loose** | FN | 1359 |

| | Predicted Label | |
|---|---|---|
| **True Label** | *Positive* | *Negative* |
| *Positive* | 3657679 | 1359 |
| *Negative* | 12 | 53049191 |

| | Statistic |
|---|---|
| **Error Rate** | 0.000024 |
| **Accuracy** | 0.999976 |
| **Precision** | 0.999997 |
| **Recall** | 0.999629 |
| **Specificity** | 1.000000 |
| **F1** | 0.999813 |

**Supplementary Figure 3**. Confusion matrix for all RGI *main* annotations, with accompanying performance metrics.
